# RaFAH: A superior method for virus-host prediction

**DOI:** 10.1101/2020.09.25.313155

**Authors:** FH Coutinho, A Zaragoza-Solas, M López-Pérez, J Barylski, A Zielezinski, BE Dutilh, RA Edwards, F Rodriguez-Valera

**Affiliations:** Evolutionary Genomics Group, Departamento de Producción Vegetal y Microbiología, Universidad Miguel Hernández, Alicante, Spain; Molecular Virology Research Unit, Faculty of Biology, Adam Mickiewicz University Poznan, Uniwersytetu Poznanskiego 6, 61-614, Poznan, Poland; Department of Computational Biology, Faculty of Biology, Adam Mickiewicz University Poznan, Uniwersytetu Poznanskiego 6, 61-614, Poznan, Poland; Centre for Molecular and Biomolecular Informatics (CMBI), Radboud University medical centre / Radboud Institute for Molecular Life Sciences, Nijmegen, The Netherlands; Theoretical Biology and Bioinformatics, Science for Life, Utrecht University (UU), Utrecht, The Netherlands; Computational Sciences Research Center, San Diego State University, San Diego, USA; Moscow Institute of Physics and Technology, Dolgoprudny 141701, Russia

**Author notes:** Address correspondence to: Felipe Hernandes Coutinho, Universidad Miguel Hernández, Dpto. Producción Vegetal y Microbiología, Aptdo. 18., Ctra. Alicante-Valencia N-332, s/n, San Juan de Alicante, Alicante, Spain, Zip Code: 03550.

## Abstract

Viruses of prokaryotes are extremely abundant and diverse. Culture-independent approaches have recently shed light on the biodiversity these biological entities^1,2^. One fundamental question when trying to understand their ecological roles is: which host do they infect? To tackle this issue we developed a machine-learning approach named Random Forest Assignment of Hosts (RaFAH), based on the analysis of nearly 200,000 viral genomes. RaFAH outperformed other methods for virus-host prediction (F1-score = 0.97 at the level of phylum). RaFAH was applied to diverse datasets encompassing genomes of uncultured viruses derived from eight different biomes of medical, biotechnological, and environmental relevance, and was capable of accurately describing these viromes. This led to the discovery of 537 genomic sequences of archaeal viruses. These viruses represent previously unknown lineages and their genomes encode novel auxiliary metabolic genes, which shed light on how these viruses interfere with the host molecular machinery. RaFAH is available at https://sourceforge.net/projects/rafah/.

---

Viruses that infect Bacteria and Archaea are the most abundant and diverse biological entities on Earth. Despite their sheer abundance and broad ecological niches these entities remain elusive. Culture-independent techniques such as metagenomics^1^ have been pivotal in the effort to describe viral biodiversity. Computational approaches have been developed to assign these novel viruses to putative hosts^3^. These rely on identifying genomic signals that are indicative of a virus-host association.

First, alignment-free methods such as *k*-mer profiles assume that viruses and their hosts have similar nucleotide composition profiles. Since expression of viral genes depends on the translational machinery of the host, viruses adapt their oligonucleotide composition to that of the host they infect. This process may be driven by the adaptation of the codon usage to tRNA pool available in the host cell, exchange of the genetic material, co-evolution of regulatory sequences and/or an evasion of the host defence systems. Hence, by identifying the prokaryote genome with the highest significant similarity to a viral genome, one can assume that prokaryote genome to be the host of the virus in question. Alignment-free methods show very high recall (i.e., percentage of viral genomes assigned to a host) but usually have low precision (i.e., percentage of correct virus-host assignments among the predicted virus-host assignments), with reported scores for genus-level predictions between 33% and 64% depending on the dataset^3–5^. Similarities in *k*-mer profiles between viruses can also be used for host prediction following the same rationale^6^.

Second, there are alignment dependent approaches to assess similarity between viral and prokaryote genomes. These methods assume that genetic exchange between viral and prokaryote genomes are indicative of virus-host associations. Specific genetic fragments, although short, might be informative for this purpose, such as CRISPR spacers, exact matches and tRNAs. While longer matches such as whole genes or integrated prophages can also provide an indication of virus-host linkage^3^. Both aforementioned approaches are limited by the fact that they require the genome of the host to be present in the reference database. Alignment dependent approaches also require that detectable genetic exchange has taken place between virus and host.

Third, manual curation can be used to investigate the gene content of viral sequences on a one-by-one basis in search of specific markers genes that are indicative of the host, such as photosynthesis genes for cyanophages^7^. The manual approach may have high precision but usually the recall of such predictions is low, the procedure is extremely time-consuming and prone to human errors.

All three approaches have been used extensively in viral metagenomic studies to assign hosts to uncultured viruses^1,7–9^. An ideal tool for virus-host prediction should combine the precision of alignment-dependent methods and the recall of alignment-free approaches. Furthermore, it should be independent of host genomes so not to be limited by database completeness biases. Previous studies have shown that Random Forest decision algorithms are the ideal tool for classifying viruses according to their hosts^10^ and that protein domains can be used to achieve accurate host predictions^11,12^. Based on these findings we postulated that Random Forest classifiers could be applied to protein domain data to build a classifier that is based on identifying combinations of genes that are indicative of virus-host associations. Through this approach we were able to design RaFAH, a classifier that combined the precision of manual curation, the recall of alignment-free approaches, and the speed and flexibility of machine learning (Figure 1).

**Figure 1:**
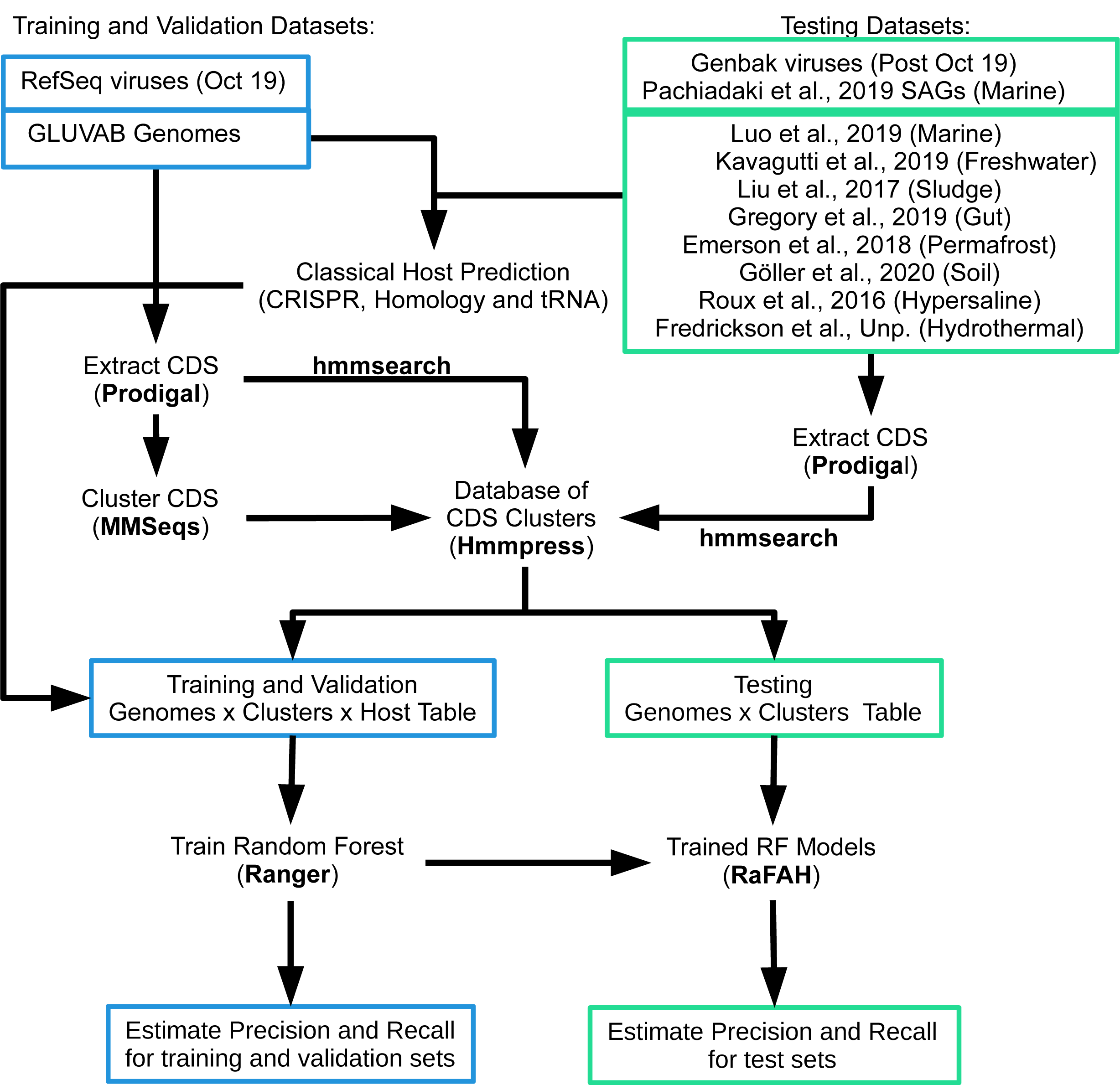
Overview of the strategy used to train, validate and test random forest models.

We tested the performance of RaFAH and other host prediction approaches on an independent dataset of isolated viral genomes that did not overlap with those used for training the models (Test Set 1). When using these methods without any cutoff (i.e. considering as valid all predictions from these approaches regardless of their score), RaFAH outperformed both alignment-independent and alignment-dependent approaches for host prediction at every taxonomic level as evidenced by the F1-score (Figure 2A). HostPhinder and CRISPRs displayed high precision only at the most strict cutoffs (Figure 2B). As a consequence, these two methods displayed very low recall when the highest cutoffs for predictions were established (Figure 2C). WiSH had a higher recall than CRISPR and HostPhinder at the expense of lower precision. Thus, the F1-score of RaFAH suggests that it can predict much more virus-host interactions than the other tested approaches while maintaining high precision, particularly for divergent viral genomes that escape detection by the classical approaches. The following RaFAH score cutoffs provided approximately 95% precision at each taxonomic level: 0 for domain, 0.14 for phylum, 0.3 for class, 0.47 for order, 0.48 for family and 0.93 for genus.

**Figure 2:**
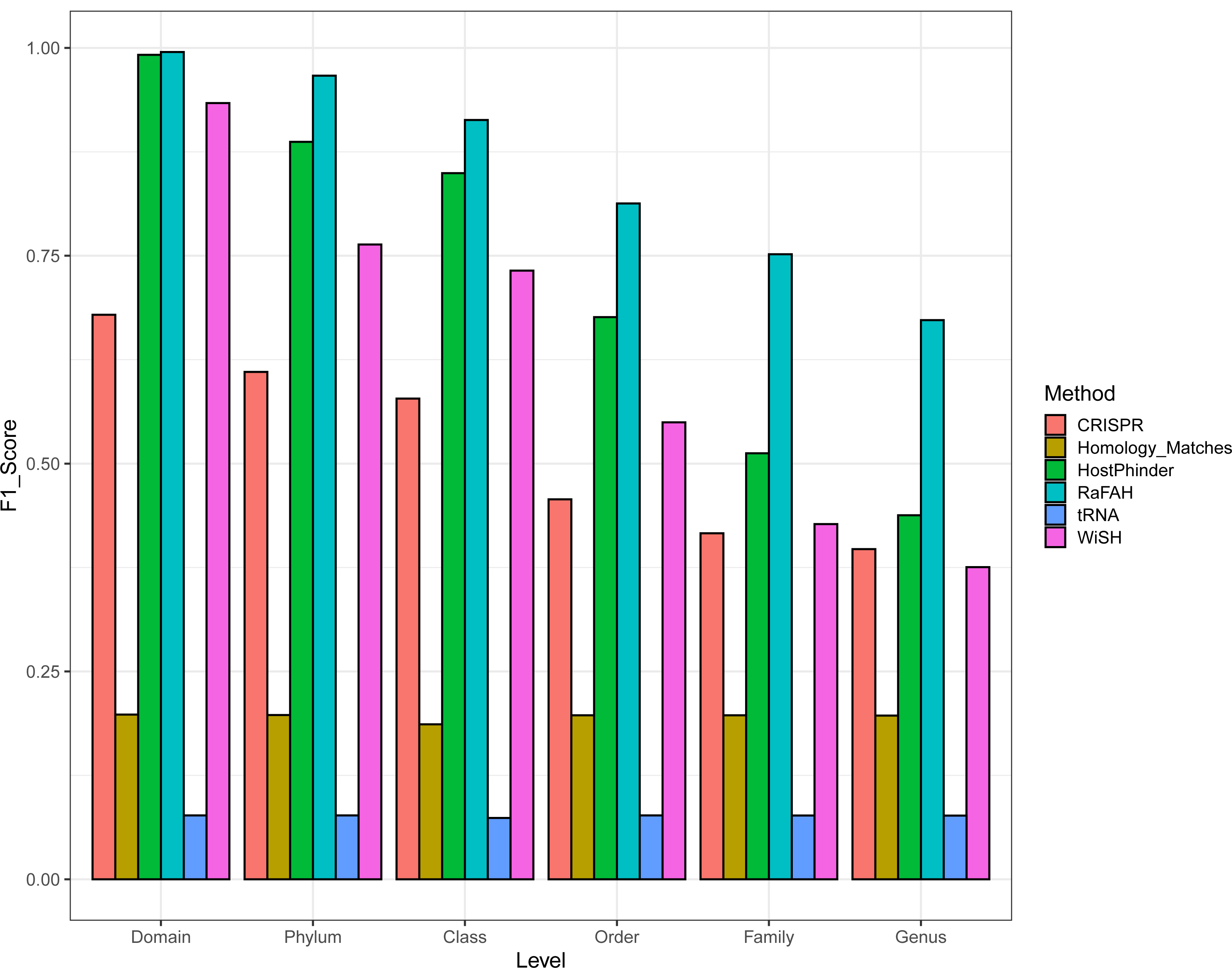
Performance of RaFAH compared to alignment-free and classical host prediction approaches on TestSet1. A) F1-score of methods when considering all predictions regardless of score at multiple taxonomic levels. B) Association between score cutoff and recall of predictions for each method. C) Association between score cutoff and precision of predictions for each method.

We applied importance analysis to determine which protein clusters were most relevant for predicting viral hosts using RaFAH. Accordingly, the most important predictor was annotated as an Rz-like phage lysis protein (Table S1). Among the protein clusters that ranked among the 50 most important ones were included multiple lysins, tail, and tail fiber proteins. These proteins are known to determine virus-host range as they play fundamental roles in virus entry and exit and host recognition^13^. The fact that these proteins ranked amongst the most important for RaFAH predictions is evidence that it learned to predict virus-host associations based on proteins which are directly involved in virus-host molecular interactions.

RaFAH was further validated on a dataset of viral genomic sequences derived from marine Single Amplified Genomes (SAGs), Test Set 2^14^. These sequences represent an ideal test dataset because they are uncultured viruses, not represented in the NCBI database used for training, and can confidently be assigned hosts because these viruses were inside or attached to the host cells during sample processing. As expected, the recall for Test Set 2 was lower than that obtained for Test Set 1, as these were divergent genomes which are more difficult to assign a putative host (Figure S2 A). Due to this lower recall, we could not confidently determine cutoffs for 95% precision for this dataset (above the 0.4 cutoff recall approaches 0 and precision stabilizes at 100%). Nevertheless, a negative association between precision and recall as a function of the score cutoff was also observed for Test Set 2 (Figure S2 C). Taken together, these results are evidence that RaFAH also performed well when predicting hosts of uncultured viruses.

To test the performance of RaFAH in other habitats we applied it to assign hosts to a dataset of viral genomes obtained from metagenomes of eight different ecosystems (Test Set 3). For comparison, we also applied the classical host prediction approaches to this dataset. When using classical approaches for host prediction the majority of viruses remain unassigned regardless of ecosystem, and the best performance of these approaches was among the human gut dataset, in which only about 25% of all the base pairs could be assigned to a host a the level of phylum (Figure 3). Meanwhile, when set to the 0.14 cutoff, RaFAH was capable of assigning putative hosts to the majority of viral sequences across all ecosystems, except for the permafrost dataset, likely because viruses derived from this ecosystem are poorly represented in reference databases.

**Figure 3:**
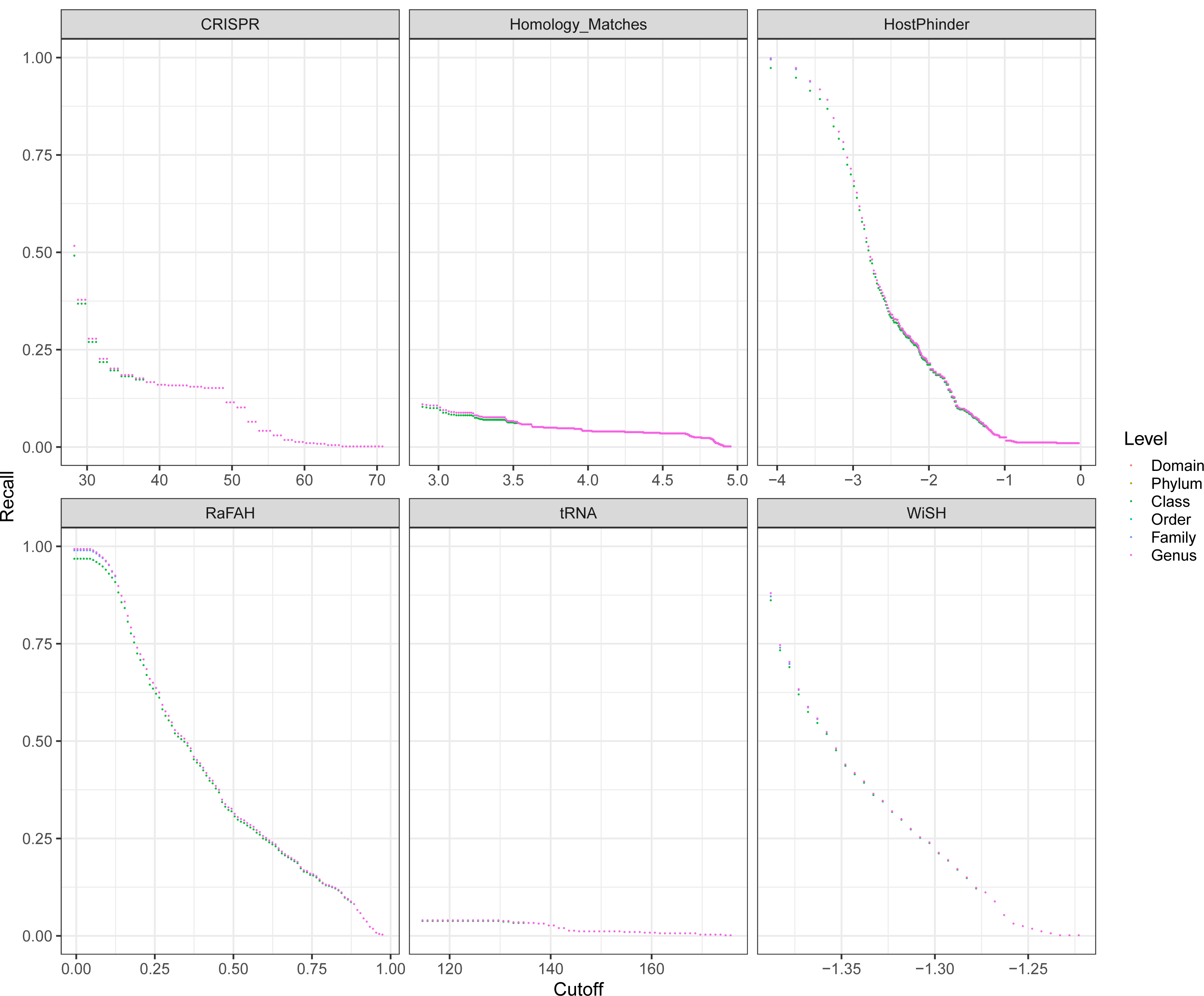
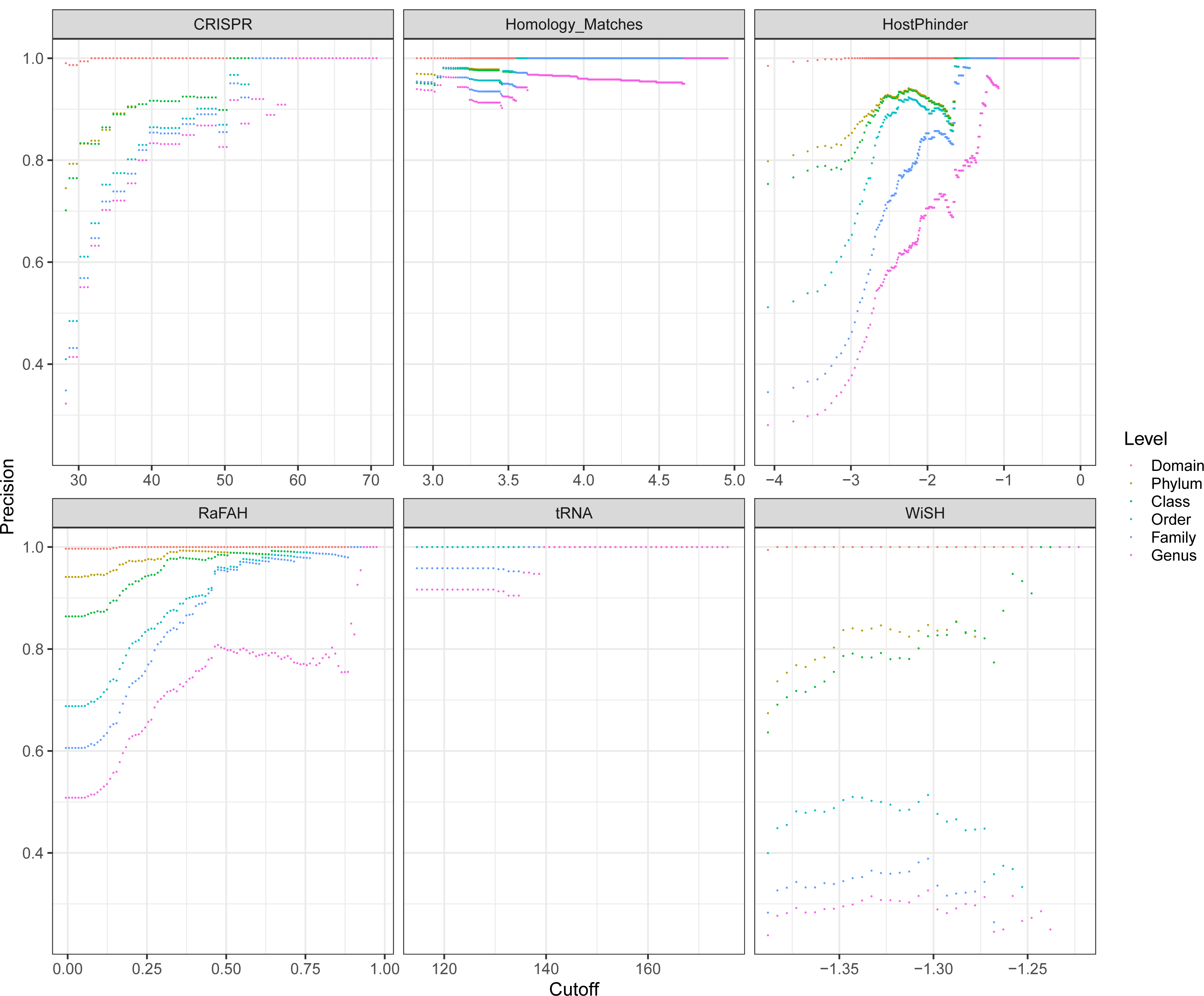
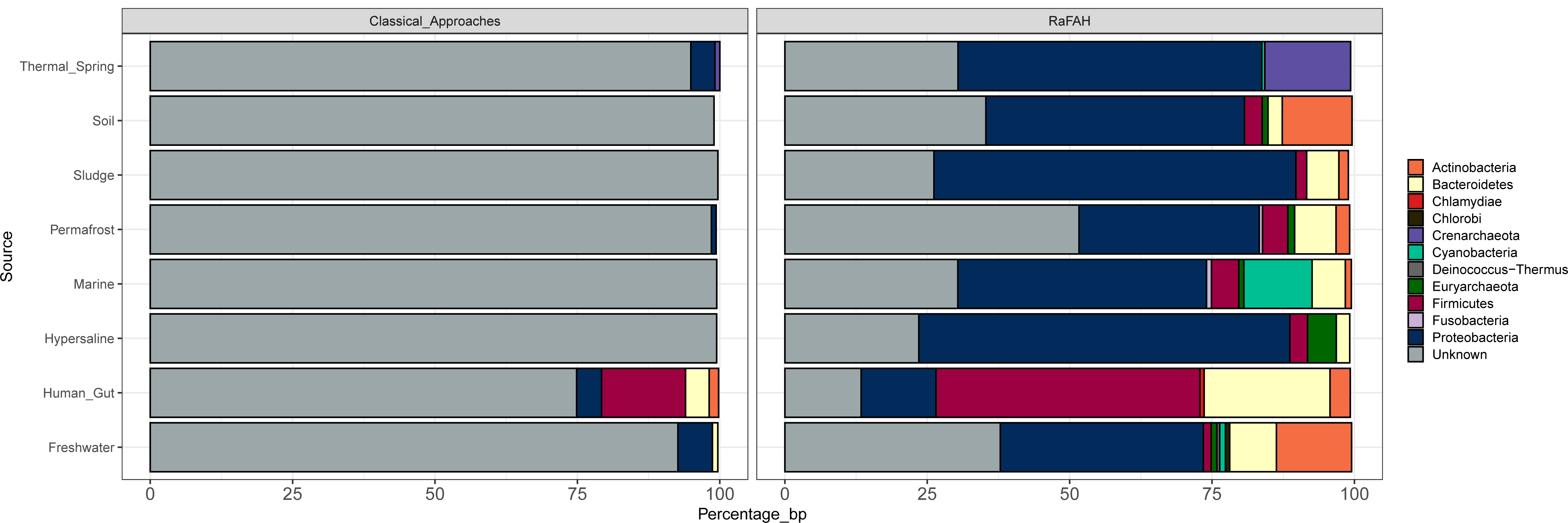
Description of the viromes of six ecosystems using classical host prediction approaches and RaFaH. For each dataset we calculated the fraction of the assembly assigned to each putative host phylum by each method.

Interestingly, the host predictions yielded by RaFAH were markedly different across ecosystems. Viruses of Proteobacteria were the dominant group in all ecosystems, except the human gut. As expected, the most abundant targeted hosts of the viruses from each ecosystem were the most abundant taxa that reside at those habitats. Viruses of Cyanobacteria were the second most abundant group among the marine dataset, a position that was occupied by viruses of Actinobacteria and Bacteroidetes among the freshwater dataset. Viruses of Firmicutes and Bacteroidetes were the dominant group among the dataset of human gut viruses. Meanwhile, viruses of Firmicutes, Bacteroidetes and Actinobacteria were among the most abundant among the soil and permafrost datasets. Viruses of Euryarchaeota were the second most abundant group among the hypersaline dataset, a position that was occupied by viruses of Crenarcheaota in the thermal springs dataset. These results are in accordance with the known prokaryote diversity that dwells in each of these ecosystems^9,15–21^. Although this agreement between virus and host community composition is to be expected, it is seldom observed in studies of viral ecology based on metagenomics, because classical methods for host prediction leave the majority of viruses unassigned. RaFAH circumvents these issues by providing accurate and complete description of viral communities regarding their targeted hosts.

Based on the finding that RaFaH achieved nearly perfect precision for Domain level host predictions, and the fact that viruses of Archaea are underrepresented in databases, we subsequently focused on the description of these viruses. Few large-scale studies have addressed the diversity of uncultured viruses of Archaea, and they focused mostly on marine samples^22–25^. Here, we describe viruses from seven other ecosystems: soil, permafrost, freshwater, sludge, hypersaline lakes, thermal springs, and the human gut. Applying RaFAH to only eight metagenomic datasets led to the discovery of 537 genomes of viruses or Archaea (prediction score ≥ 0.14). To put this figure in context, there are only 96 genomes of viruses of Archaea deposited in the NCBI RefSeq database.

We took several steps to ensure that these genomes were truly derived from viruses of Archaea and consistently found compelling evidence to our claim. First, these genomes could be linked to archaeal genomes either through homology matches or alignment independent approaches, which provided further evidence that 423 out of the 537 genomes (79%) were derived from archaeal viruses (Table S2). Second, much like the RefSeq genomes of archaeal viruses, these sequences were enriched in Pfam domains annotated as exclusive of archaea, eukaryotes and their viruses (Figure S3). Third, these genomes were enriched in ribosomal binding site motifs that are also enriched among RefSeq viruses of archaea (Figure S4).

Next, we manually inspected the gene content of the viruses predicted to infect Archaea in search for novel auxiliary metabolic genes (AMGs) and new mechanisms of interaction with the host molecular machinery. The small number of reference genomes of Archaea and their viruses makes it difficult to describe the gene content of the archaeal viruses that we discovered because most of their genes have no taxonomic or functional annotation. However, we have found several sequences containing genes coding for thermosomes, group II chaperonins involved in the correct folding of proteins, homologous to their bacterial counterparts, GroEL/GroES^26^. Other AMGs found among archaeal viruses, were those involved in the synthesis of cobalamin *cob*S, recently associated with Marine Group I (MGI) Thaumarchaeota virus infection^25^ as well as genes that encoded 7-cyano-7-deazaguanine synthase QueC involved in archaeosine tRNA modification^27^. One of the AMGs most prevalent among archaeal viral genomes encoded for a molybdopterin biosynthesis MoeB protein (ThiF family). This family of proteins is involved in the first of the three steps that make up the ubiquitination process^28^. This system regulates several cellular processes through post-translational modification of proteins such as their function, location, and degradation making it an ideal target from the point of view of viruses to facilitate their replication^29^.

In conclusion, we developed a new virus-host prediction tool with great potential for studies of viral biodiversity and ecology. RaFAH outperformed all other methods that we tested and displayed high accuracy and recall in a dataset of cultured viruses, which extended to uncultured viruses from a diverse set of ecosystems. By analysing a small set of ecosystems RaFAH allowed for a significant expansion of the archaeal virosphere and shed light on their yet poorly understood content of auxiliary metabolic genes. Future studies will describe even more uncultured viral sequences and RaFAH will likely play a role on describing their hosts and allowing us to decipher their ecological roles.

## Methods

### Viral genomes database for model training and validation

Two datasets of viral genomes were used for both training and validating the random forest models. The first dataset contained the genomes of viruses of Bacteria and Archaea from NCBI RefSeq available on October 2019, which comprised 2,668 genomes along with their associated host data. To avoid overestimating precision due to identical and nearly identical genomes in the database, this dataset was made non-reduntant using CD-Hit ^30^ at a clustering cutoff of 95% identity over 50% alignment of the shorter sequence. The second dataset was comprised of the 195,698 GLUVAB genomes ^31^. GLUVAB is a database of uncultured viral genomes compiled from multiple studies that covered several ecosystems. Only those sequences classified as *bona fide* viruses of prokaryotes in the original publication were used in subsequent analysis.

### Classical host prediction for GLUVAB genomes

To use GLUVAB genomes for training and validation of the random forest models we first had to assign them to putative hosts using classical approaches. To minimize errors during this step we opted for using only alignment dependent methods due to their higher precision ^3^. The RefSeq genomes of Bacteria and Archaea were used as the reference database, We used three lines of evidence for virus-host associations: CRISPR spacers, homology matches, and shared tRNAs. CRISPR spacers were identified in the RefSeq genomes as previously described ^32^. The obtained spacers were queried against the sequences of *bona fide* viral sequences using BLASTn v2.6.0+ (task blastn-short). The cut-offs defined for these searches were: minimum identity of 100%, minimum query coverage of 100%, with no mismatches and maximum e-value of 1. Homology matches were performed by querying viral sequences against the databases of prokaryote genomes using BLASTn ^33^. The cut-offs defined for these searches were: minimum alignment length of 500 bp, minimum identity of 95% and maximum e-value 0.001. tRNAs were identified in viral scaffolds using tRNAScan-SE v1.2 ^34^ using the bacterial models. The obtained viral tRNAs were queried against the RefSeq database of prokaryote genomes using BLASTn. The cut-offs defined for these searches were: minimum alignment length of 60 bp, minimum identity of 97%, minimum query coverage of 95%, maximum of 10 mismatches and maximum e-value of 0.001. These steps for host assignment did not include the prophages in the GLUVAB database, as we were already confident of their host assignments.

We developed a per viral population scoring method. First, all GLUVAB genomes were clustered into viral populations (VPs) on the basis of 95% average nucleotide identity and 80% shared genes ^35^. For each virus-taxon association signal detected (i.e. homology, tRNA or CRISPR) 3 points were added to the taxon if it was a CRISPR match, 2 points if it was a homology match and 1 point if it was a shared tRNA. The taxon that displayed the highest score was defined as the host of the viral population. With this approach we ensured that all the genomes in the same VP were assigned to the same host and that no sequences had to be excluded due to ambiguous predictions.

### Protein cluster inference and annotation

Protein sequences were identified in viral genomes using Prodigal ^36^ in metagenomic mode. Hidden Markov Models (HMMs) for the phage proteins were built as follows: The 4,701,074 identified proteins were clustered by the cluster workflow of the MMseqs2 software suite ^37^, with parameters: 35% sequence identity and alignment coverage had to cover at least 70% of both proteins. Protein clusters (PCs) were aligned into multiple sequence alignments (MSAs) using QuickProbs ^38^ with default parameters, then converted into HMMs using the hmmake program from the HMMER suite ^39^, which resulted in 144,613 HMMs. The HMM profiles were annotated by performing HMM-to-HMM annotation against the pVOG database ^40^ using the HH-suite3 software suite ^41^. First, the MSAs provided on the pVOGs website and the ones built in the previous step were converted into the hhsuite proprietary HMM format using hhmake. The pVOG HHMs were built into a hh-suite3 database, which was then used to find matches to the phage protein HMMs using hhsearch. All HMMs could be annotated through this approach but only 4,578 matches displayed target coverage ≥ 50% and e-value ≤ 1^-10^.

Finally, individual viral proteins were mapped to the HMM profiles using the hmmsearch program limiting hits to those with i-evalue ≤ 10^−5^, alignment length ≥ 70% for both proteins, and minimum score of 50. These results were parsed into a matrix of viral genomes x PCs. Once the matrix of genomes x PC was defined we calculated Pearson correlation coefficients (*r*) between all possible pairwise combinations of PC. To remove redundancies, PCs were grouped into superclusters if they presented *r* ≥ 0.9 and only a single PC from each supercluster was kept for subsequent analysis. This reduced table of genomes versus PC scores (25,884 genomes x 43,644 PCs) was used as input to train, validate and test the random forest models.

### Random Forests training, validation and testing

Our rationale was that the machine could learn the associations between genes and hosts much more efficiently than a human, while also using the information contained in the hypothetical proteins. Hence, random forest models were built using the Ranger^42^ package in R^43^. The response variable was the genus level host assignment of the viral sequences while the input parameters were the scores of viral genomes to each PC. Random forests were built with 1,000 trees, 5,000 variables to possibly split at in each node, and using probabilistic mode. Variable importance was estimated using the impurity method. Three models were built and validated on independent datasets. Model1 was trained on Training Set 1, which comprised 80% randomly selected non-redundant viral genomes from NCBI RefSeq. The performance of this model was evaluated on Training Set 1 and Validation Set 1, which comprised the remaining 20% of non-redundant RefSeq genomes. This process was repeated for a 10-fold cross-validation. Model2 was trained on Training Set 2 which comprised 100% of the RefSeq genomes and validated on Validation Set 2 which was comprised of GLUVAB genomes that could be assigned to a host at the level of genus by the pipeline described above. Finally, Model3 was built based on Training Set 3, which was comprised of all the RefSeq viral genomes and the GLUVAB genomes that could be assigned to a host at the level of genus (i.e. a combination of TrainingSet2 and ValidationSet2). Models 1 and 2 were used as a proof-of-principle models and Model3 was the definitive model used for testing and which is provided to the users and used for all subsequent analyses.

We used three independent test sets to evaluate the performance of RaFAH Model3. Test Set 1 was comprised of viral genomes retrieved from NCBI Genbank database. We took several steps to make sure that Test Set 1 represented a challenging dataset for the random forest model, so as to assess its ability to extrapolate. First, we excluded from Test Set 1 any genomes made public in or before October 2019. Second, Test Set 1 was made non-redundant at 95% nucleotide identity and 50% alignment length of the shorter sequence. Third, protein sequences derived from Test Set 1 were compared the protein sequences of the Training Set 3 using DIAMOND ^44^. Any genomes that shared more than 80% of proteins or more than 80% average amino acid identity with any genome from Training Set 3 were removed from Test Set 1. These steps resulted in an independent Test Set 1 consisting of 600 (out of the initial 3,199) genomes with no overlap to the genomes used to train the models.

Test Set 2 was comprised of viral genomes identified in single amplified genomes (SAGs) from marine samples ^14^. A total of 4,751 SAGs (with completeness ≥ 50% and contamination ≤ 5% as estimated by CheckM^45^) were classified at the level of genus using BAT ^46^. This algorithm assigns taxonomic affiliations to microbial genomes based on consesus taxa of proteins matches to the NCBI-nr database. Next, viral sequences were extracted from the SAGs using VIBRANT ^47^ which identified 418 viral sequences. We assumed that the viral sequences in the SAGs infected the organisms from which these SAGs were derived, either because they were derived from integrated prophages or from viral particles attached or inside host cells. Viral sequences for which the host taxon predicted by RaFAH was the same taxon assigned to the SAG by BAT were considered as correct host predictions. Viruses from SAGs that could not be classified were excluded from the precision and recall analyses.

Test Set 3 was comprised of a collection of 61,647 viral genomic sequences from studies that spanned multiple samples from permafrost ^9^, marine ^48^, human gut ^49^, freshwater ^18^, soil ^50^, hypersaline lakes ^51^ and hydrothermal springs (Fredrickson et al., unpublished data obtained from IMG/VR ^52^) and a sludge bioreactor ^17^ habitats. These sequences were assigned to putative hosts through the classical host prediction pipeline described above for the GLUVAB genomes and also using RaFAH.

### Comparison with other methods for host prediction

To assess the performance of RaFAH compared to other host prediction tools we assessed the performance of the alignment free methods HostPhinder ^6^ and WiSH ^4^ and the alignment-dependent approaches based on homology matches, shared tRNAs and CRISPR spacers. For this comparison we used Test Set 1 as it was comprised of sequences from cultured viruses with known hosts. Both HostPhinder and WiSH were run with default parameters. The classical host predictions (CRISPR, tRNA and homology matches) for Test Set 1 were performed using the same parameters described above for the GLUVAB genomes and Test Set 3.

For the approaches that required a database of reference host genomes (i.e. WiSH, CRISPR, tRNA and homology matches) the database of host genomes was the NCBI RefSeq genomes of Bacteria and Archaea and the genomes of Uncultured Bacteria and Archaea (UBA) from GTDB ^53^. To minimize false-positives due to homology between viruses and mobile genetic elements all sequences that matched the keyword “plasmid” in their description field were removed from the database of reference host genomes.

### Assessment of archaeal virus host predictions

To confirm the assignment of 537 phages predicted by RaFAH as archaeal viruses we used Mash v. 2.1 ^54^. Mash calculates Jaccard distance between two genomes based on the number of shared *k*-mers with a certain length. We used *k*-mer sizes from 13 to 20 nucleotides. For each *k*-mer size we calculated distances of every phage genomic sequence against all potential host genomes. This database included 17,134 bacterial genomes and 4,716 archaeal genomes retrieved from RefSeq and GenBank. For each phage genome, we selected the potential host with the smallest Mash distance. In addition to Mash distance, we also calculated Manhattan distances and correlation scores between phage and host *k*-mer frequencies using *k* = 6 as described in ^3,5^. Finally, all 537 phages were used as BLASTn queries against the whole NR database. For each phage we assigned potential host by selecting the top-scoring non-viral hit as described in ^3^. In addition, we compared the prevalence of ribosomal binding site (RBS) motifs (defined by Prodigal ^36^ gene predictions) between viral sequences assigned to bacteria and archaea, both from the eight metagenomic datasets and RefSeq viruses. A similar analysis was performed to compare the prevalence of Pfam domains among these groups. For this analysis, protein sequences were queried against the Pfam database using hmmsearch with maximum e-value set to 10^−3^.

## Data availability

All the data (Viral and Prokaryote genomes) analyzed in this study is freely available from public repositories.

## Code availability

RaFAH and the associated files necessary to run it are freely available online at https://sourceforge.net/projects/rafah/

## Acknowledgments

This work was supported by grants “VIREVO” CGL2016-76273-P [MCI/AEI/FEDER, EU] (cofounded with FEDER funds) from the Spanish Ministerio de Ciencia e Innovación and “HIDRAS3” PROMETEU/2019/009 from Generalitat Valenciana. FRV was also a beneficiary of the 5top100-program of the Ministry for Science and Education of Russia. FHC was supported by APOSTD/2018/186 post-doctoral fellowships from Generalitat Valenciana. AZ was funded by Polish National Science Centre [2018/31/D/NZ2/00108]. JB research was supported by the National Centre for Research and Development (NCBR, Poland), grant number LIDER/5/0023/L-10/18/NCBR/2019. BED was supported by the Netherlands Organization for Scientific Research (NWO) Vidi grant 864.14.004 and by the European Research Council (ERC) Consolidator grant 865694: DiversiPHI.

## Author contributions

FHC conceived and designed the experiments. FHC, AZS, MLP, AZ, JB, BED, and RAE analyzed the data. All authors contributed to writing the manuscript.

## Competing interests

The authors declare that they have no competing interests.

## Ethics Approval and Consent to Participate

Not Applicable.

## Consent for publication

Not Applicable.

## Supplementary material

Figure S1: Associations between score cutoff, precision, and recall for RaFAH, the alignment-free (WiSH and HostPhinder), and classical host prediction approaches (CRISPR, tRNA and homology matches) on TestSet1.

Figure S2: Associations between score cutoff, precision, and recall for RaFAH on Test Set 2.

Figure S3: Prevalence of Pfam domains among viruses. Pfam domains were grouped according to their expected taxonomic ranges (depicted above each panel). Only values derived from scaffolds with at least 5 CDS are shown to reduce noise. A) Comparisons of Pfam domain prevalence between RefSeq viruses of Archaea and Bacteria. The *p*-values of each comparison obtained with the Mann-Whitney test are depicted above bars. B) Comparisons Pfam domain prevalence between RefSeq viruses of Archaea and Bacteria, viruses from TestSet3, and uncultured viruses from previous publications reported as viruses of Archaea. Notice the different y axes on each panel.

Figure S4: Prevalence of ribosomal binding site (RBS) motifs among viruses. Only values derived from scaffolds with at least 5 CDS are shown to reduce noise. A) Comparisons of RBS motif prevalence between RefSeq viruses of Archaea and Bacteria. The *p*-values of each comparison obtained with the Mann-Whitney test are depicted above bars. B) Comparisons RBS motif prevalence between RefSeq viruses of Archaea and Bacteria, viruses from TestSet3, and uncultured viruses from previous publications reported as viruses of Archaea.

Table S1: Variable importance and protein cluster annotation derived from Model3. Importance of protein clusters for RaFAH predictions was estimated based on the impurity method. Protein clusters queried against the pVOGs database for functional annotation.

Table S2: Details regarding putative viruses of Archaea identified by RaFAH. Table includes sequence identifies, source dataset, sequence length, score of host prediction performed by RaFAH, host taxonomy (Domain to Genus), and results from the alignment dependent (BLASTN) and alignment independent (Mash and *k*-mer distances) best matches to RefSeq genomes of Archaea.

## Notes

### Competing Interest Statement

The authors have declared no competing interest.

