## Supplementary figures and images for "RaFAH: A superior method for virus-host prediction"

### Figure S1

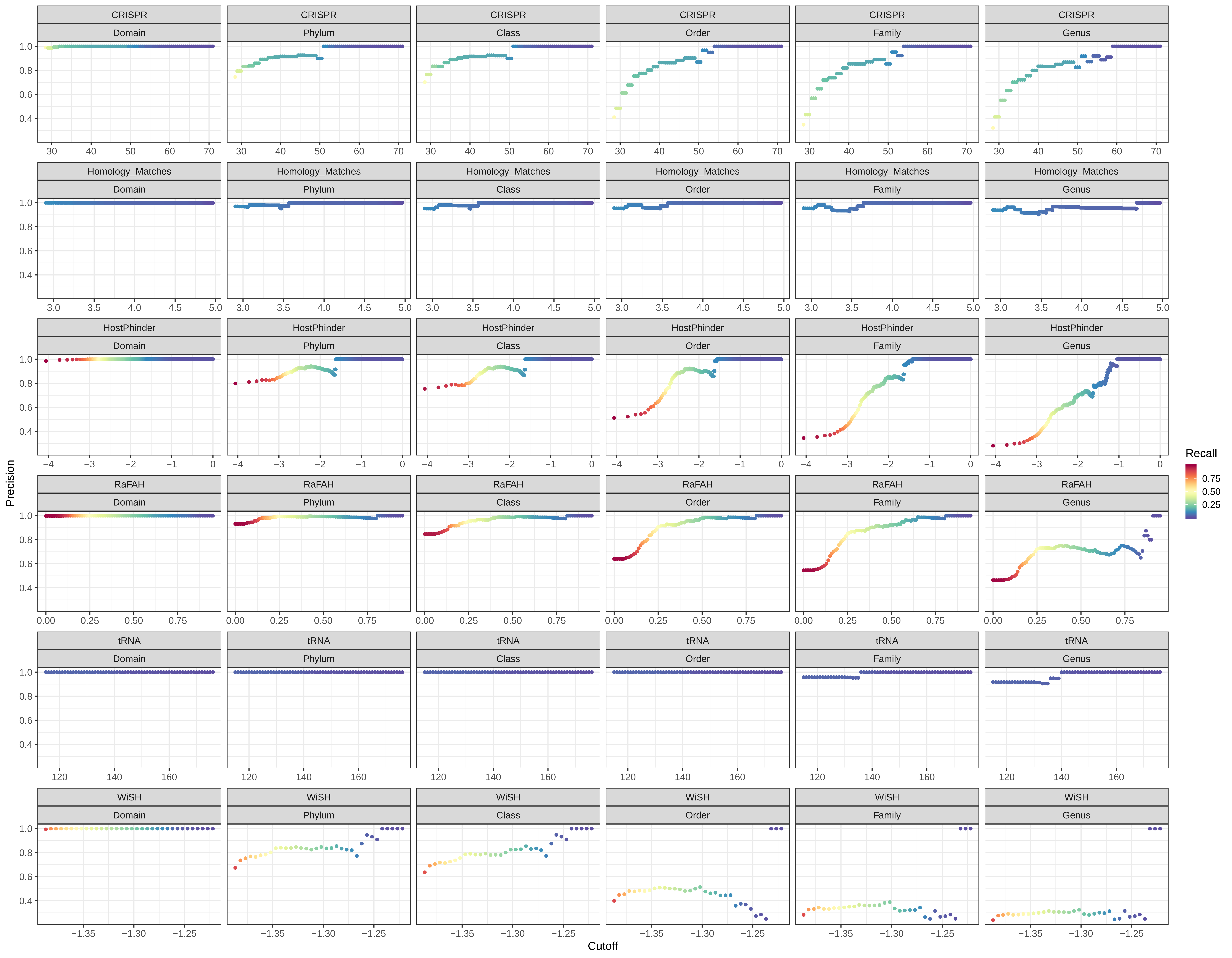

### Figure S2 A

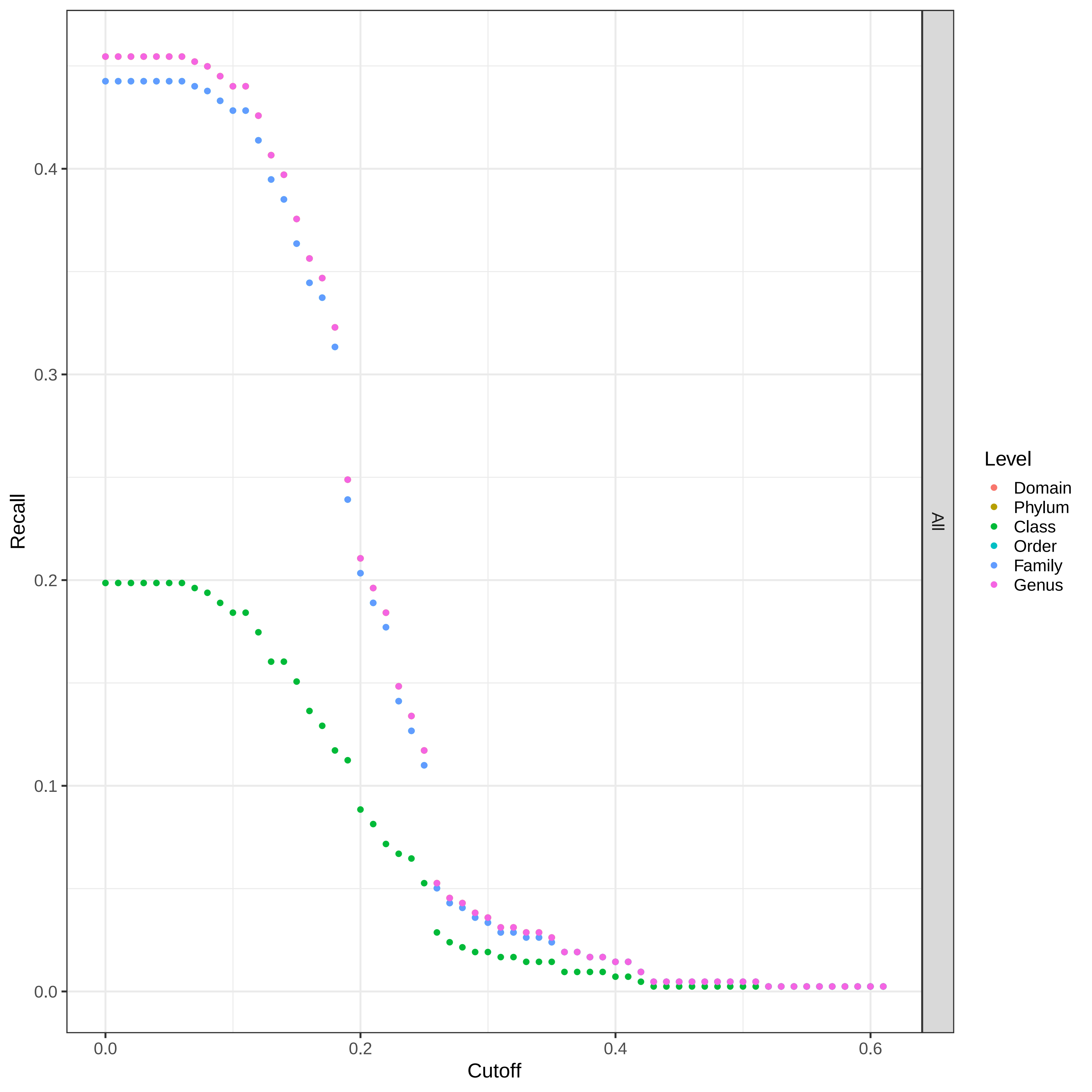

### Figure S2 B

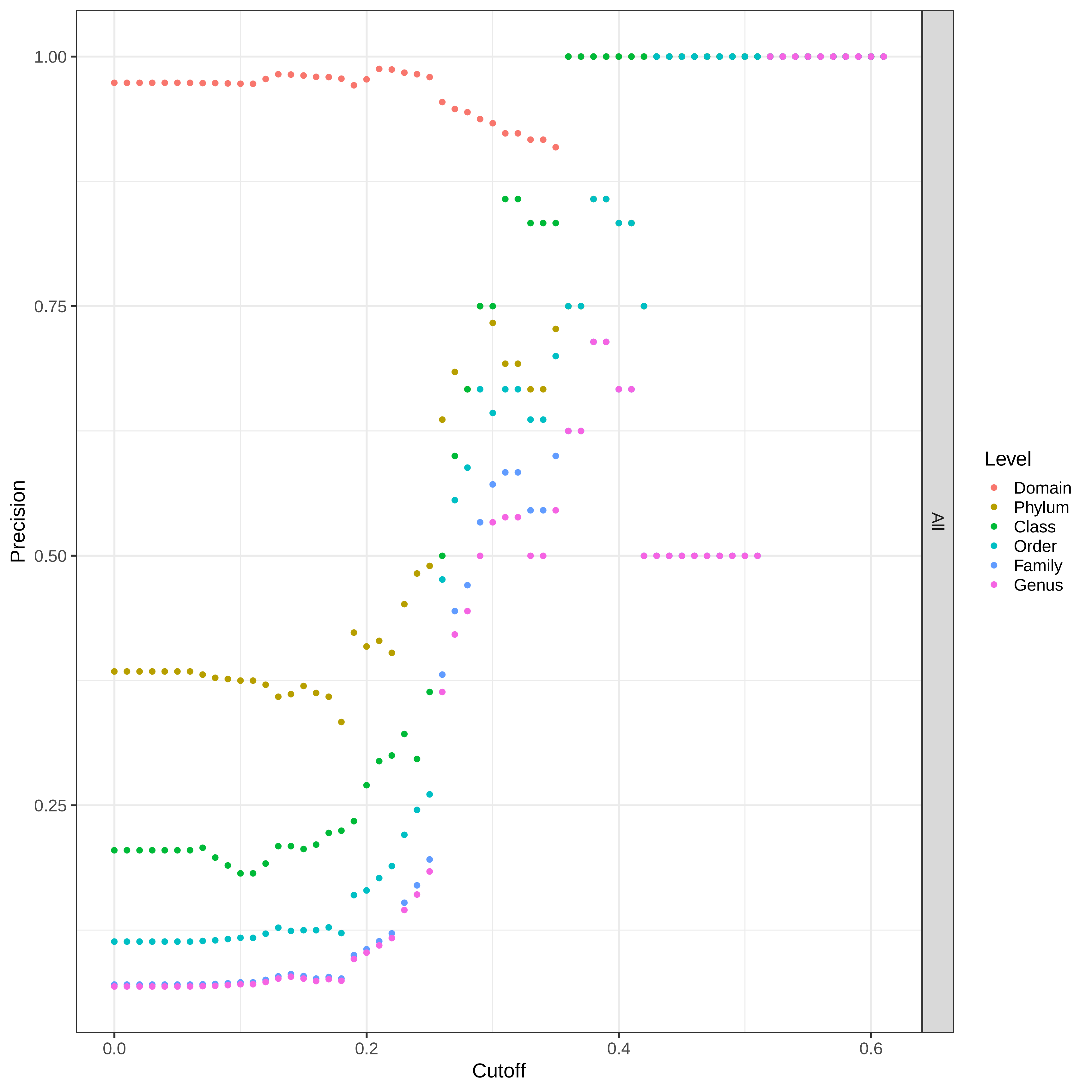

### Figure S2 C

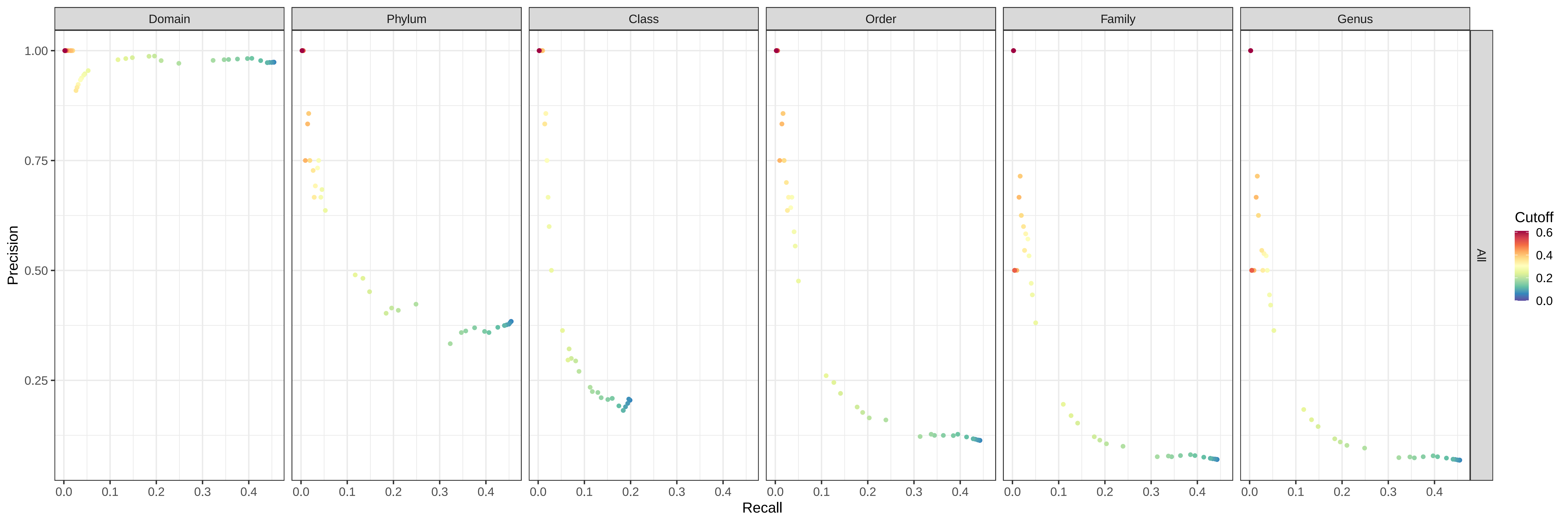

### Figure S3 A

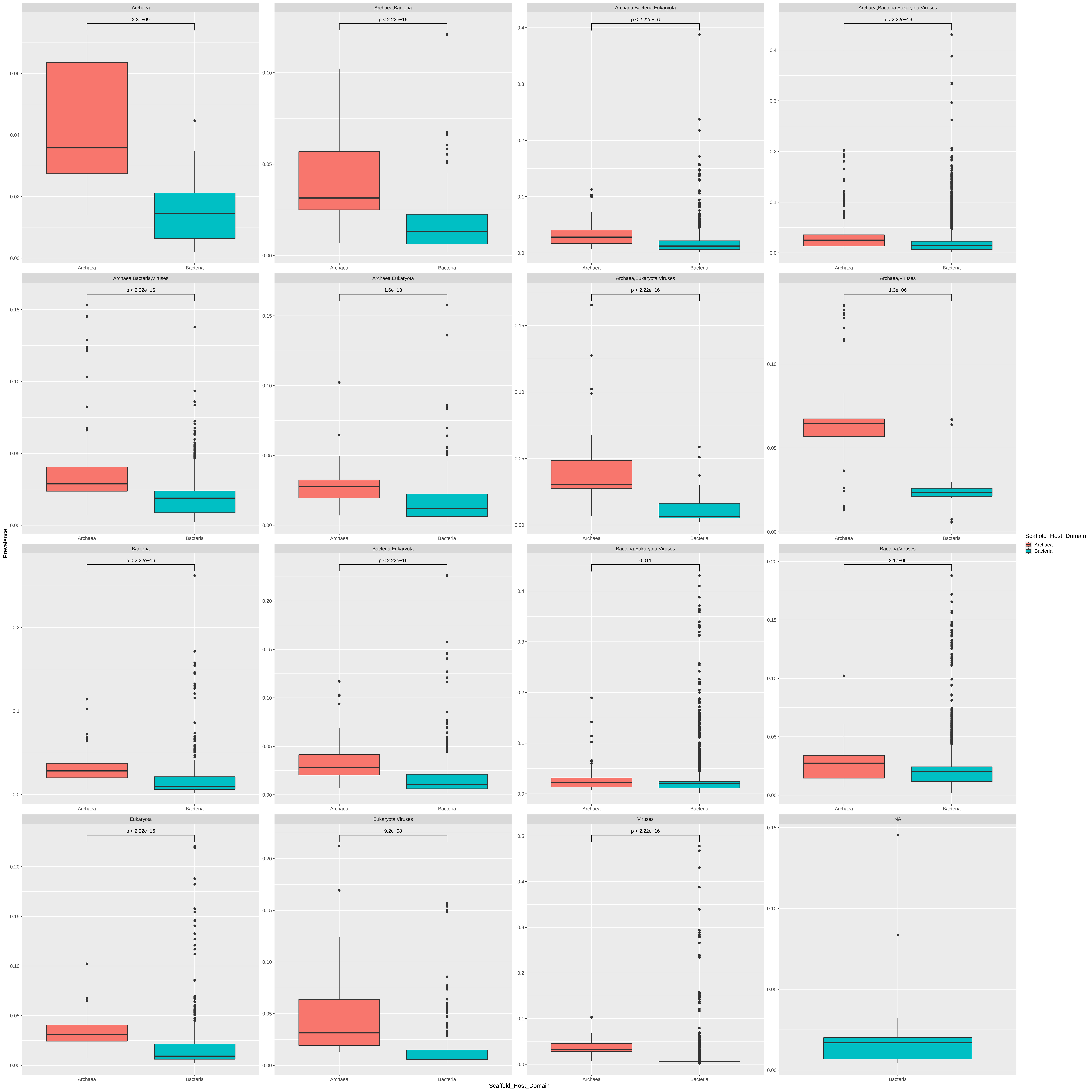

### Figure S3 B

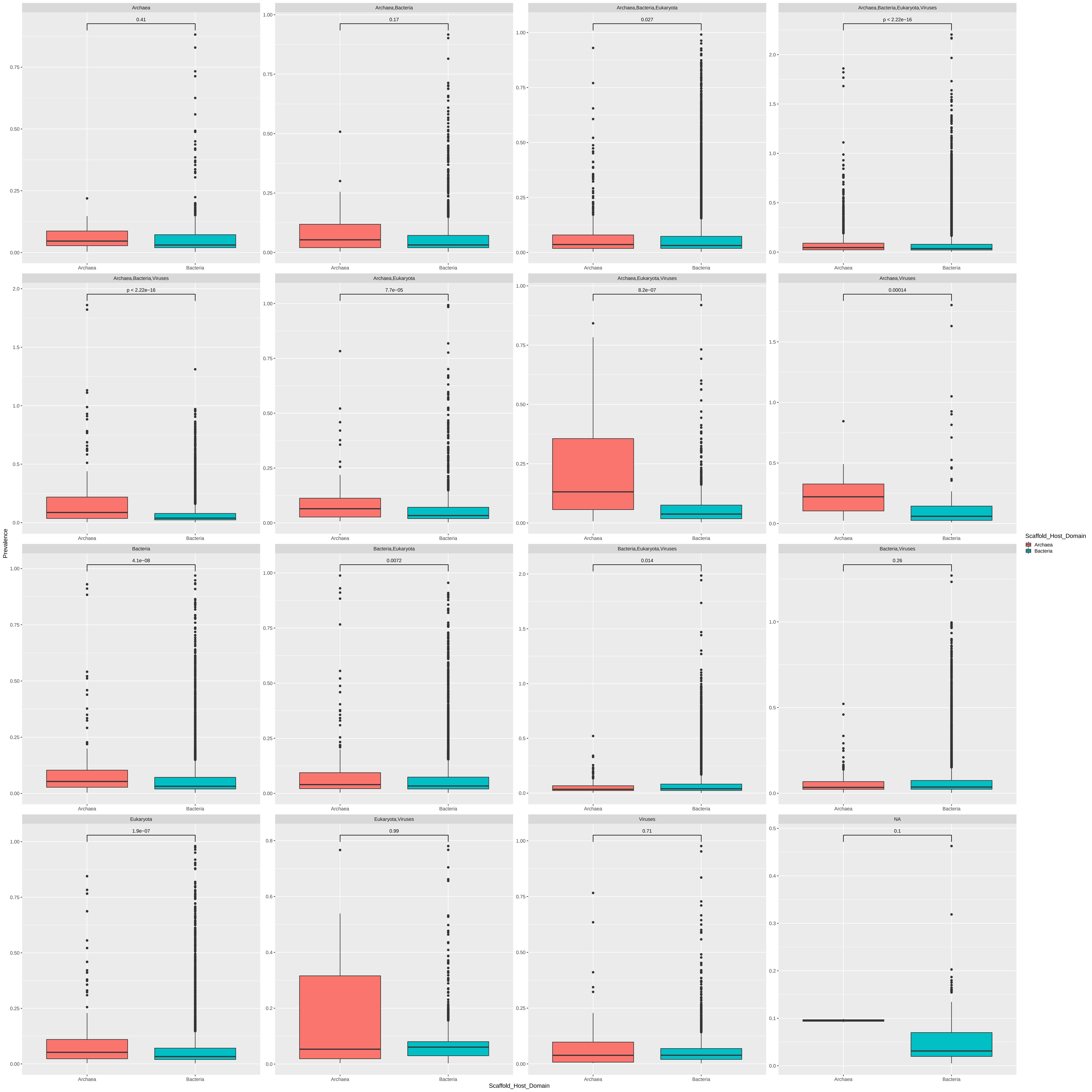

### Figure S4 A

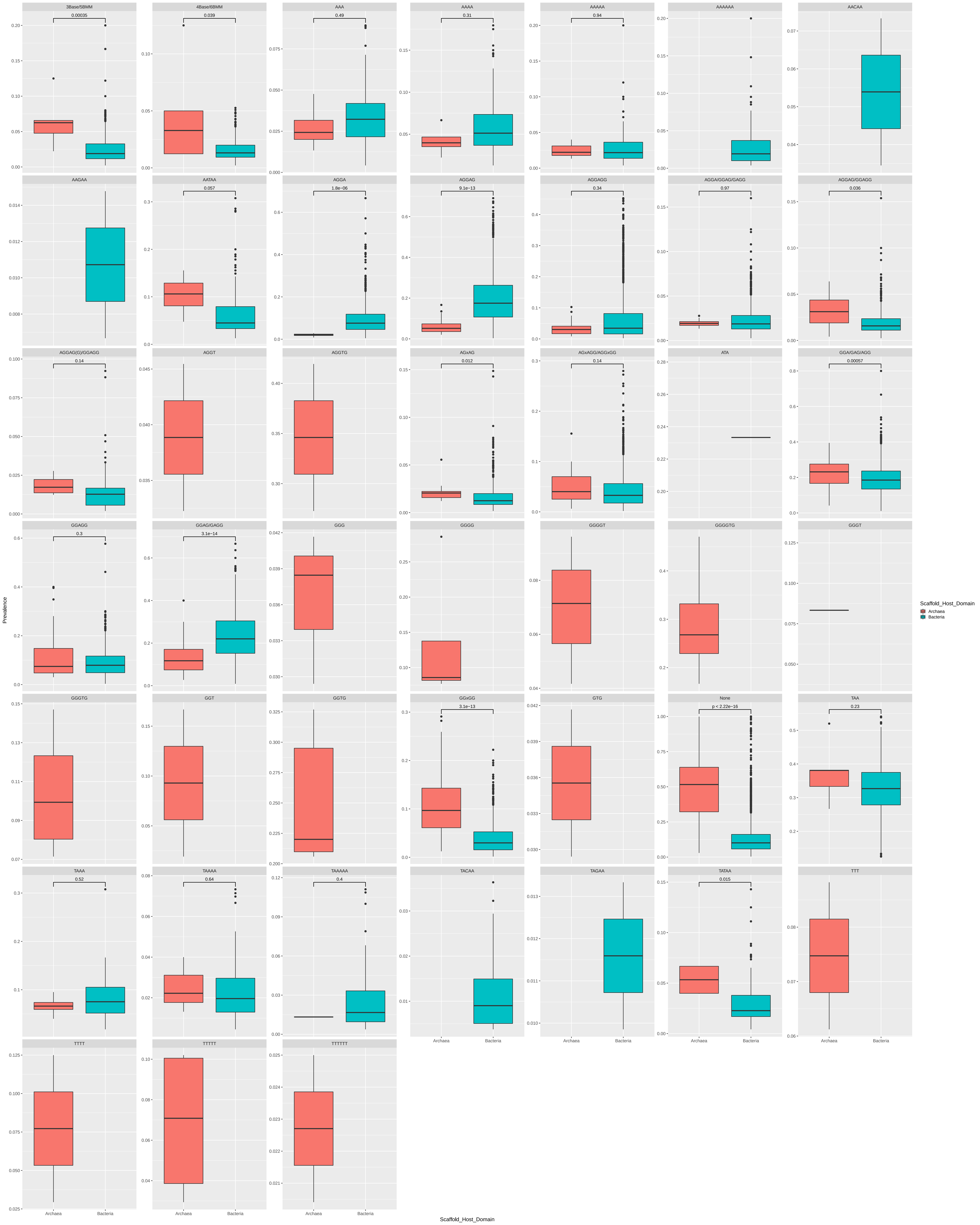

### Figure S4B

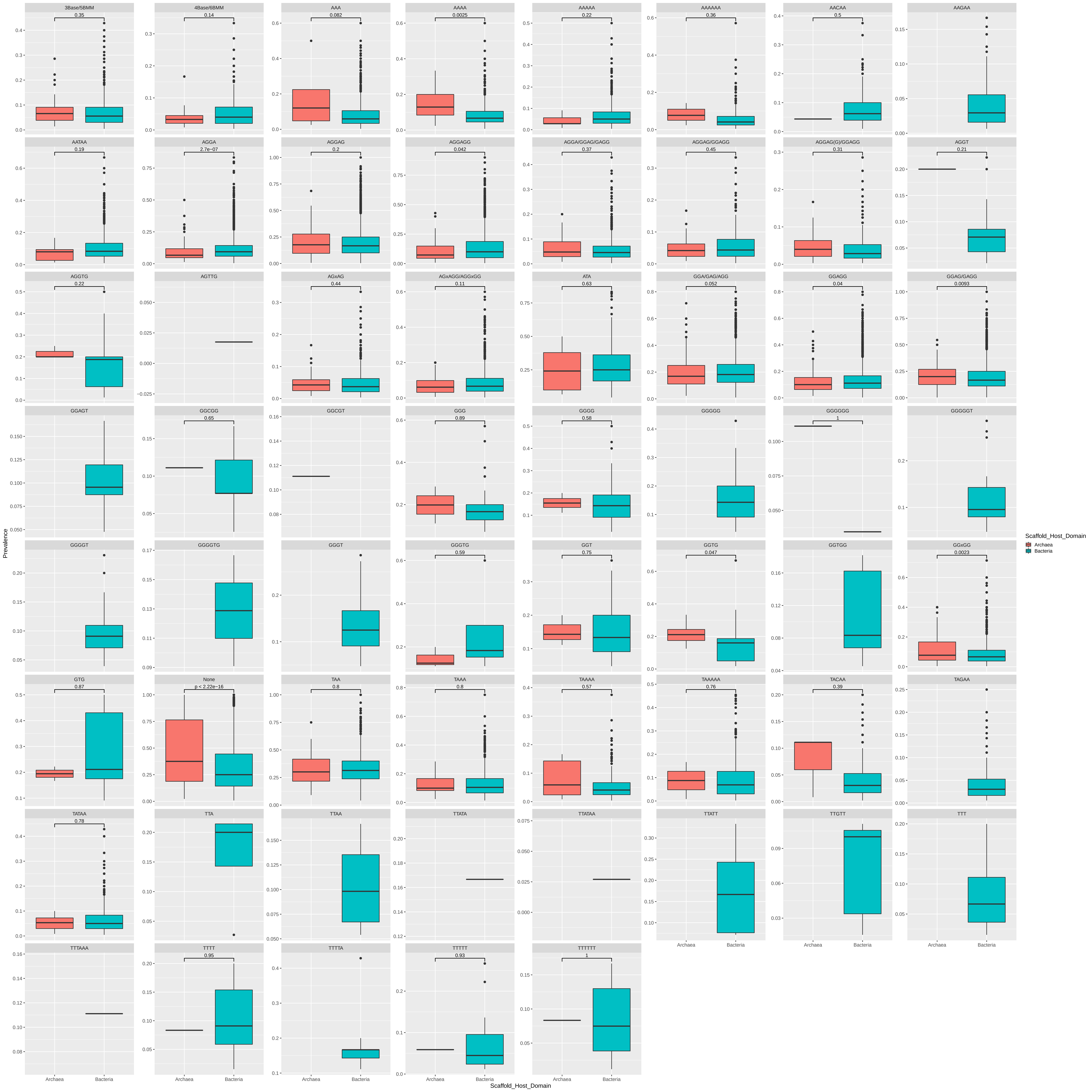
